# Motility of Enzyme-Powered Vesicles

**DOI:** 10.1101/645986

**Authors:** Subhadip Ghosh, Farzad Mohajerani, Seoyoung Son, Darrell Velegol, Peter J. Butler, Ayusman Sen

## Abstract

Autonomous nanovehicles powered by energy derived from chemical catalysis have potential applications as active delivery agents. For *in vivo* applications, it is necessary that the engine and its fuel, as well as the chassis itself, be biocompatible. Enzyme molecules have been shown to generate mechanical force through substrate turnover and are attractive candidates as engines; phospholipid vesicles are biocompatible and can serve as cargo containers. Herein, we describe the autonomous movement of vesicles with membrane-bound enzymes in the presence of the substrate. We find that the motility of the vesicles increases with increasing enzymatic turnover rate. The enhanced diffusion of these enzyme-powered systems was further substantiated in real time by tracking the motion of the vesicles using optical microscopy. The membrane-bound protocells that move by transducing chemical energy into mechanical motion serve as models for motile living cells and are key to the elucidation of the fundamental mechanisms governing active membrane dynamics and cellular movement.

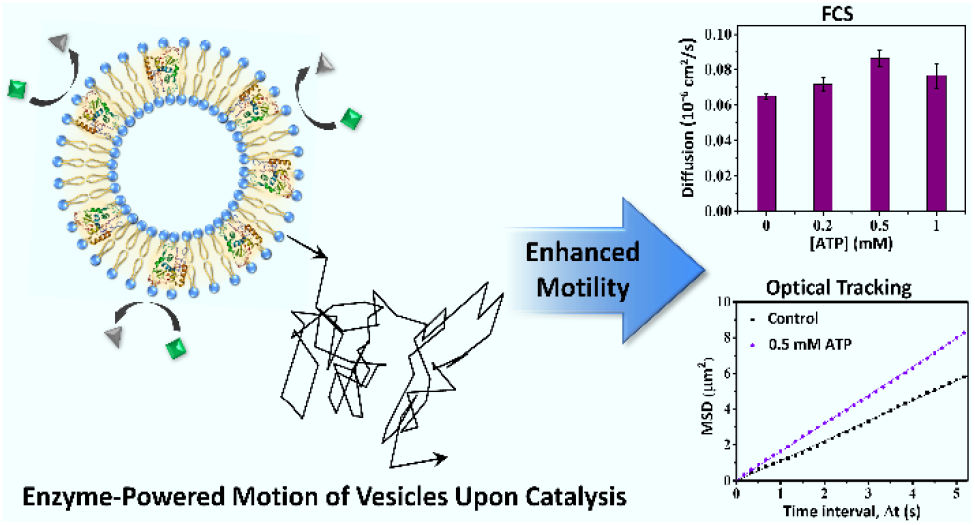
“For Table of Contents Only”

## Introduction

The diffusive movement of enzymes have been shown to increase significantly during substrate turnover, suggesting force generation through catalysis.^1–7^ In principle, the phenomenon can be exploited to power nanovehicles by harnessing enzymes to the vehicle chassis.^3,8^ The combination of biocompatible enzymatic engine and a biocompatible cargo container allows the design of delivery vehicles that can be employed *in vivo*.^9–12^ Herein, we show that phospholipid vesicles with embedded enzymes exhibit enhanced mobility in presence of the substrate. In addition to employing free-swimming enzymes such as acid phosphatase (AP) and urease, we have also examined the behavior of adenosine 5’-triphosphatase (ATPase), a member of the membrane-bound enzyme family. These enzymes span across the phospholipid bilayer of the cell membrane and regulate cellular activity by controlling transport of ions and molecules across the membrane.^13–16^ Following observations on free swimming enzymes, we anticipated that ATPase will also exhibit self-generated fluctuations. These dynamic fluctuations may lead to the fluctuations in the membrane itself or even enhanced motion of the entire cellular structure (Figure 1A); a time dependent phenomenon that remains virtually unexplored.

**Figure 1:**
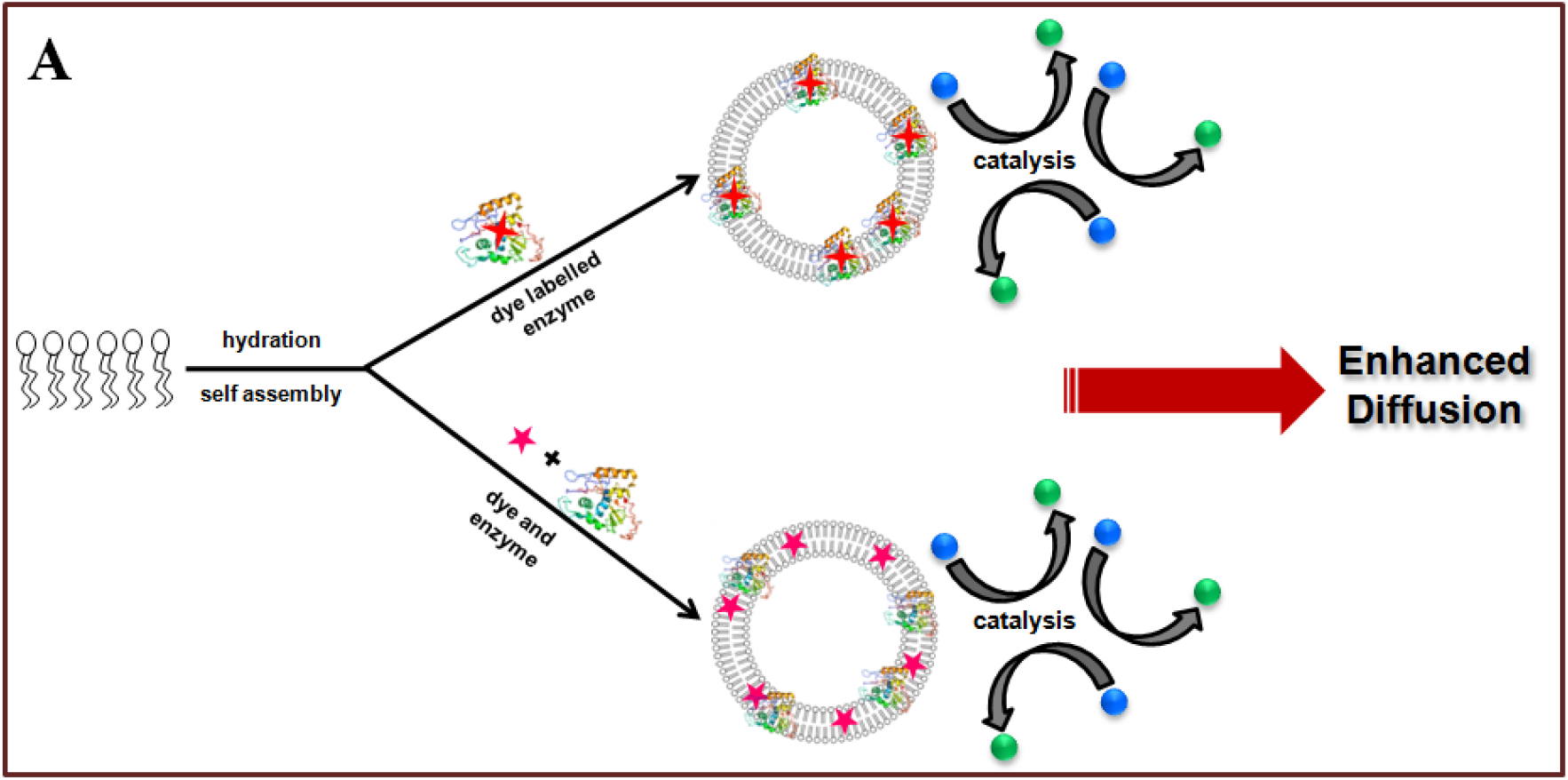
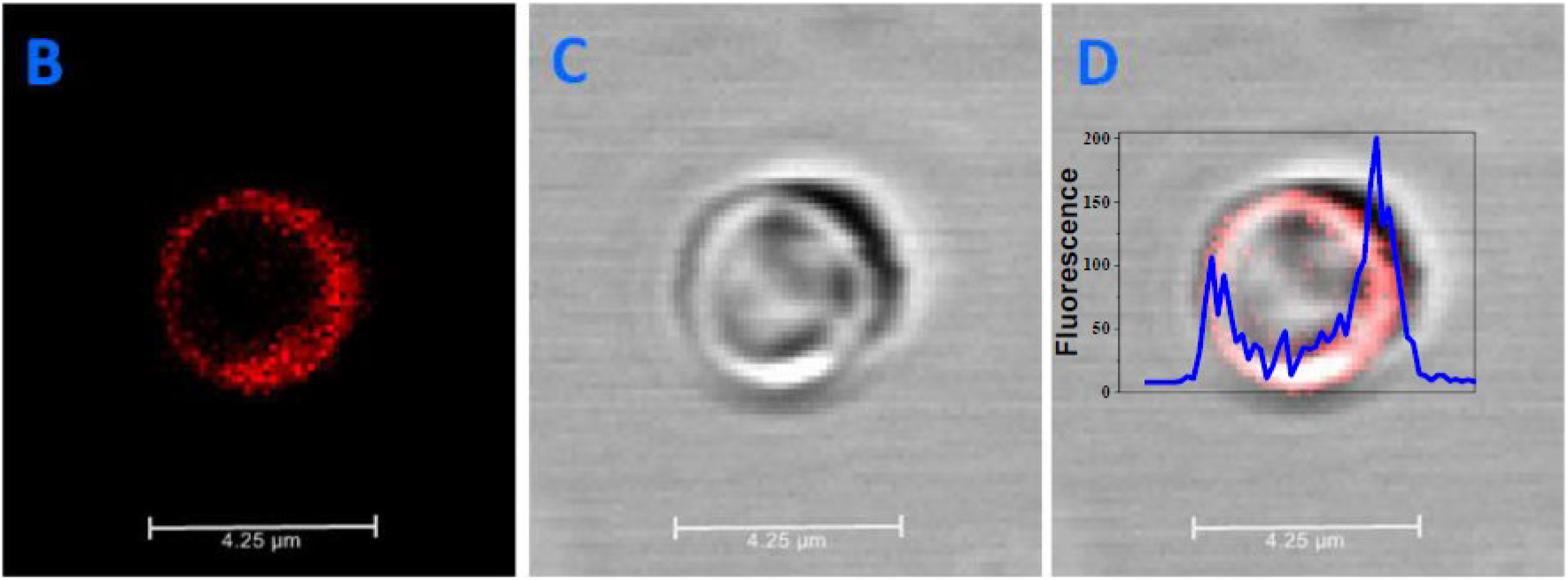
(A) Schematic representation showing the enhanced diffusion of enzyme tagged vesicle due to substrate turnover. Confocal image of a single large vesicle tagged with Chromeo P540 labelled ATPase. (B) Fluorescence channel (C) Bright field image (D) Merged image. The fluorescence intensity profile along *z*-axis scan is overlaid with the confocal image of the vesicle.

A phospholipid lipid vesicle was chosen which can provide the necessary parameters to study enzyme-powered diffusion.^10^ Fluorescence correlation spectroscopy (FCS) was used to probe the diffusion changes of a conventional membrane-bound enzyme, Na^+^/K^+^ activated ATPase after it was reconstituted in the vesicle bilayer. In addition, to examine the generality of enzyme-catalyzed enhanced mobility of vesicles, we studied two other enzymes, AP and urease which were covalently attached to the vesicle membrane through biotin-streptavidin conjugation. The three-dimensional enhanced diffusion of these enzyme-powered systems was further substantiated in real time by tracking the motion of the vesicles using optical microscopy.

## Results and Discussion

The principal enzyme in this study, Na^+^/K^+^ ATPase, is a well-known transmembrane (integral) enzyme which typically regulates the cell membrane potential by controlling the concentration gradients of Na^+^ and K^+^ ions across the membrane.^17^ The energy it acquires from catalyzing the conversion of adenosine triphosphate (ATP) to adenosine diphosphate (ADP) is utilized to pump Na^+^ and K^+^ ions in opposite directions (Figure S1).^18^ The diffusion behavior of free Na^+^/K^+^ ATPase was first studied in solution by FCS, in the presence of varying concentrations of ATP. For conducting FCS measurements, ATPase was fluorescently labelled with Chromeo P540 dye. The pyrylium dye fluoresces only on conjugation with the primary amines.^19^ The dye covalently reacts and selectively labels ATPase. After tagging, the diffusion coefficient of a single ATPase was measured at nanomolar enzyme concentrations. On fitting the autocorrelation curves, the following results were obtained: in the absence of ATP, the diffusion coefficient of ATPase was 0.19 × 10^−6^ cm^2^/s. Upon addition of ATP to ATPase solution in the presence of 1 mM Mg^+^ ions and at fixed pH, the diffusion coefficient of the enzyme increases to a maximum value of 0.26 × 10^−6^ cm^2^/s with 0.5 mM ATP (Figure S2). Thus, an enhancement of ∼37% in diffusion was observed when compared to the diffusion value of ATPase without ATP. At higher concentration of 1 mM ATP, ATPase diffusion coefficient shows a drop, presumably due to product inhibition (Figure S2).

Transmembrane enzymes usually aggregate in aqueous medium and hence purification strategies are important to separate out non-aggregated, active enzymes. It has also been suggested that ATPase may dissociate into its subunits in dilute solutions.^20^ Even though diffusion changes were clearly observed for free ATPase in solution, it is preferable to construct the natural environment for the transmembrane ATPase by assembling it within lipid membranes. The vesicle synthesis procedure in brief involved vacuum drying of L-α-phosphatidylcholine (EPC) dissolved in chloroform, to form a lipid film. The subsequent rehydration step with ATPase solution ensures reconstitution of the enzyme within the bilayer following reported protocols with minor modifications (Figure S3, details in the Supporting Information).^21–24^ To impart fluorescent characteristics to the vesicle, either the ATPase was fluorescently labelled with Chromeo P540 dye or the lipid molecules were labelled with Nile red (NR) dye. The free ATPase and dye molecules were removed from the solution by dialysis. Scanning electron microscopy was performed to confirm the formation of vesicles (Figure S4A). In addition, confocal microscopy scans were also carried out for fluorescent vesicles. Both methods confirm formation of lipid vesicles (Figure S4). The confocal scan of a single large vesicle tagged with Chromeo P540 labelled ATPase molecules show strong fluorescence along the vesicle boundary as indicated by “horn like” fluorescence intensity profile (Figure 1B-D). This image supports the successful reconstitution of ATPase molecules within the vesicle membrane during its synthesis.

FCS, which was primarily used in this work for observing the changes, is a very sensitive technique for measuring the diffusion changes of molecules.^25–28^ The autocorrelation curves for vesicle tagged and free ATPase show distinct differences in the diffusion time **τ**_D_, which is directly related to the size of the system according to Stokes-Einstein equation (Figure S5). To prepare homogeneous and uniform sized ATPase tagged vesicles for FCS measurements, the vesicles were extruded through polycarbonate filters. The resulting solution of vesicles obtained after 50 passes, have vesicle sizes ∼100 nm. The Nanosight particle analyzer plot shows a sharp peak at 100 nm suggesting size uniformity (Figure S6). The enzymes tagged vesicles are termed as ‘active’ whereas the vesicles without enzymes are ‘inactive’. For FCS experiments, the ATPase tagged vesicle solution was diluted to nanomolar concentration to ensure that diffusion changes originate from a single vesicle in the sample volume. The Chromeo P540 labelled ATPase tagged vesicles exhibited a change in the diffusion coefficient from 0.072 × 10^−6^ cm^2^/s in the absence of ATP, to a highest value of 0.094 × 10^−6^ cm^2^/s on the addition of 0.5 mM ATP; the difference corresponds to an enhancement of ∼31% in diffusion (Figure 2A). From the FCS measurements, we also obtained the value for the average number of ATPase per vesicle of 4.5 ± 0.9. Alternatively, the dye labelled ATPase was replaced with NR labelled lipids to synthesize vesicles with fluorescently labelled membrane. ATPase was reconstituted into NR labelled vesicles to examine whether the position of the dye has any effect on the diffusion of the active vesicles. The diffusion plots from both the experiments are essentially identical, with an estimated increase of ∼32% observed for NR labelled vesicles. The increase was evaluated from the diffusion coefficient of 0.065 × 10^−6^ cm^2^/s in buffer and 0.086 × 10^−6^ cm^2^/s in the presence of 0.5 mM ATP (Figure 2B).

**Figure 2:**
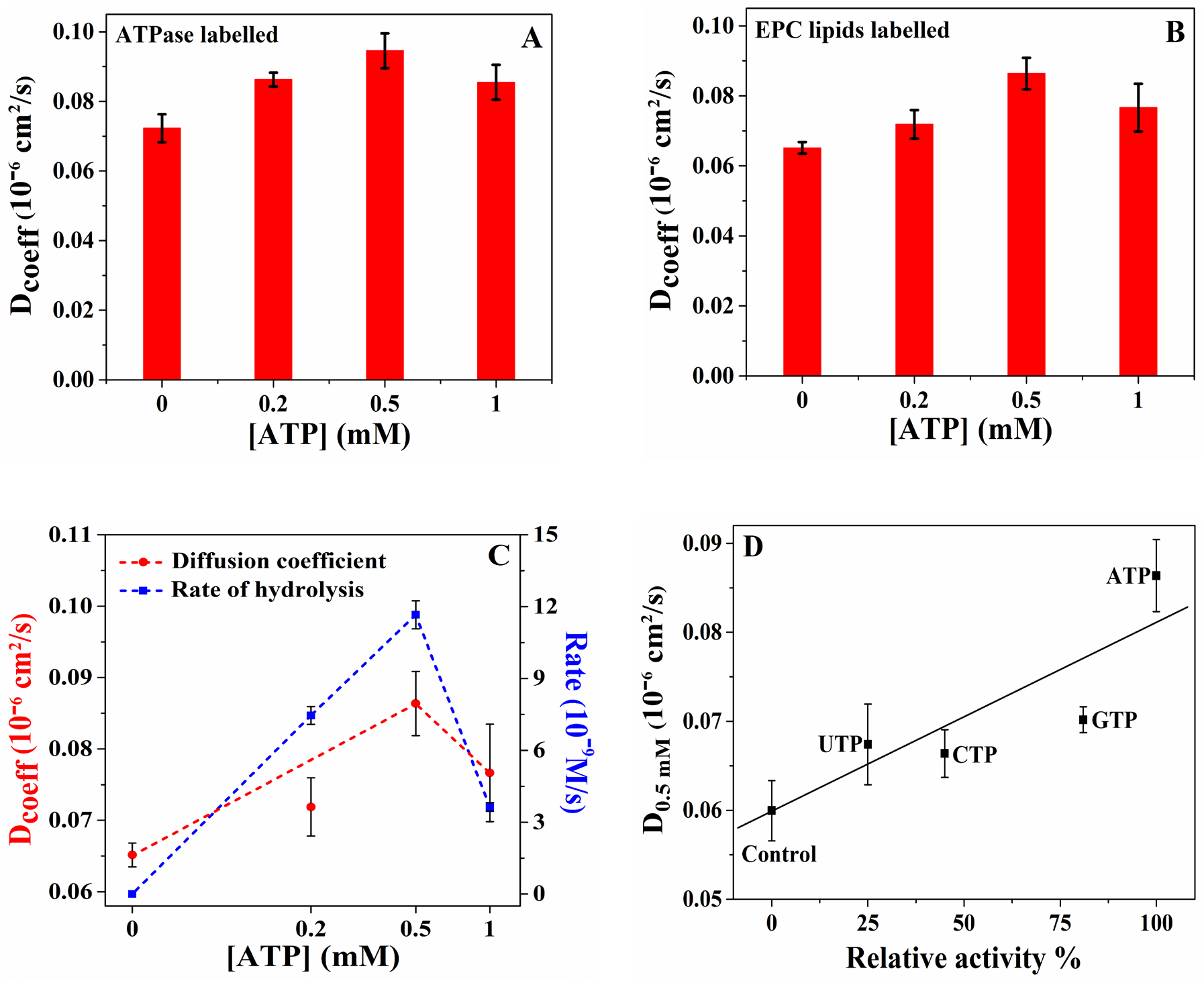
Diffusion coefficient (D_coeff_): (A) Chromeo P540 labelled ATPase tagged to vesicles and (B) ATPase tagged to Nile Red labelled lipid vesicles, in the presence of Mg^2+^ and increasing ATP concentration. (C) Overlay plot showing the rate of ATP hydrolysis with ATPase tagged vesicles (blue) versus the diffusion data from FCS (red) against ATP concentrations along the x-axis. (D) The plot of diffusion of ATPase tagged vesicles in the presence of 0.5 mM of different trinucleotides versus their % activity relative to ATP. The relative activity % was calculated from the reported k_cat_ values of the trinucleotides.^29^ Conditions for all the measurements: [Mg^2+^] = 1 mM; [HEPES buffer] = 10 mM (pH = 7.8). Diffusion plots were obtained from the average of five FCS recordings. All the FCS experiments were performed at T=294 K ± 1 K.

Applying the same FCS procedure, two separate control experiments were performed with inactive vesicles instead of active ones to determine the effect on its diffusion due to 1) the activity of free ATPase in solution, and 2) the presence of substrate and/or ions. In the first control experiment, the diffusion of NR labelled inactive vesicles was monitored in the presence of 10 nM fixed concentration of free ATPase while varying the ATP concentration. In the second control, diffusion of NR labelled inactive vesicles was measured in the presence of fixed MgCl_2_ concentration and varying concentrations of ATP. In both control experiments, the diffusion coefficient of the vesicles remained essentially unchanged (Figure S7A,B). The above results suggest that catalysis by ATPase reconstituted in the vesicles is solely responsible for the observed enhancement in the diffusion of ATPase tagged vesicles in the presence of ATP. The enzymatic catalysis is accompanied by the conversion of chemical energy into mechanical force causing enhanced diffusion of the active vesicles.^3,8^

The drop in the diffusion of ATPase tagged vesicles at high ATP concentrations was also scrutinized. FCS analyses were performed with active vesicles in the presence of the reaction products, ADP and sodium phosphate. Although the diffusion of the vesicles remained unchanged with Na_3_PO_4_, the diffusion changes of the active vesicles with ADP was observed (Figure S8). This suggests that ADP may remain bound to ATPase and inhibit ATP hydrolysis at high ADP concentrations. To further probe the relationship between the rate of ATP hydrolysis and the diffusion enhancement for ATPase, we performed an assay to estimate the reaction rate at different ATP concentrations. Malachite green (MG) dye was used as the probe for estimating the amount of phosphate ion produced from the hydrolysis of ATP by ATPase. Buffered solutions of ATPase tagged vesicles were titrated with varying ATP concentrations and the reaction mixture was added to the acidified MG solution in fixed time intervals (details in the Supporting Information). The typical reaction is: H_3_PMo_12_O_40_ + HMG^2+^→ (MG^+^)(H_2_PMo_12_O_40_) + 2H^+^. The corresponding UV peak due to the formation of green colorimetric complex (MG^+^)(H_2_PMo_12_O_40_) was monitored at 630 nm.^30^ As shown in Figure 2C, the diffusion of ATPase tagged vesicles is strongly correlated with the ATP hydrolysis rate. As seen, both drop at high ATP concentrations.^31,32^ We also examined whether the diffusion of ATPase tracks the known hydrolysis rate for different triphosphates by this enzyme. FCS experiments were carried out for ATPase tagged vesicles with guanosine triphosphate (GTP), cytidine triphosphate (CTP) and uridine triphosphate (UTP). As shown, when the value of the diffusion coefficient at 0.5 mM for each triphosphate was plotted against the enzyme activity relative to ATP, an approximate correlation was observed (Figure 2D).

To establish that the observed catalysis-induced enhanced diffusion of active vesicles is not limited to transmembrane enzymes, the diffusion behavior of two other enzymes, AP and urease, was explored. The enzymes were chemically attached to the vesicle membrane using biotin-streptavidin (BS) linkage. The BS linkage is one of the strongest known chemical bonds (K_d_≈ 10^−14^ M) and thus it is expected that BS bonds attach the enzymes firmly on the vesicle membrane.^33^ The protocol ensures that diffusion changes are only due to the attached enzyme on the vesicle surface which basically mimic the ‘peripheral membrane enzymes’.^16^ A solution of the enzyme was functionalized initially with biotin molecule; subsequently after purification, the biotinylated enzyme solution was allowed to react with streptavidin (details in the Supporting Information). The lipid used contained a biotin functionality attached to the hydrophilic head group. Hence the addition of BS-tagged solution of the enzyme to the biotinylated lipid yielded vesicles that are covalently tagged with the enzyme (Figure 3A). NR was used to fluorescently label the lipid molecules prior to the lipid drying step. Similar to the protocol used for the FCS measurements of ATPase, AP/urease linked vesicle solutions were extruded and then diluted to nanomolar concentration prior to the FCS measurements. AP is an enzyme that catalyzes the hydrolysis of organic phosphates at acidic pH.^34^ Prior to the study of vesicle bound AP, the diffusion of free AP was measured by varying the concentration of substrate, 4-nitrophenyl phosphate (PNPP). AP was labelled with AlexaFluor532 succinimidyl ester and diluted to nanomolar concentration before experiments. Free AP in buffer exhibited a diffusion coefficient of 0.71 × 10^−6^ cm^2^/s. On the addition of 1 mM PNPP, the diffusion coefficient shows maxima of 1.01 × 10^−6^ cm^2^/s, a 42% enhancement (Figure S9A). The diffusion coefficient shows saturation after 1 mM of PNPP. On the other hand, the AP tagged vesicle exhibited a diffusion coefficient of 0.057 × 10^−6^ cm^2^/s in the absence of PNPP. The diffusion coefficient increased gradually with PNPP attaining a highest value of 0.071 × 10^−6^ cm^2^/s at 1 mM of PNPP (Figure S9B). The increase in diffusion coefficient for AP tagged vesicles was estimated to be 25%. In this context, it is important to mention that PNPP and other nitro compounds are reported to be fluorescence quenchers which can drastically reduce the intensity and lifetime of the fluorophore in its close vicinity.^20,35^ Fluorescence lifetime measurements for the free AP versus PNPP concentrations was thus performed; the corresponding curve fitting confirmed significant drop in lifetime of the Alexa dye used for labelling AP (Figure S10A). Hence, as an alternative, experiments were done by replacing PNPP with ATP as substrate. The lifetime values of the Alexa dye remained unchanged with varying ATP concentrations (Figure S10B) signifying that ATP does not affect the lifetime and quantum yield of the dye. Hence the diffusion measurements of AP using FCS are free from artefacts under these conditions. The diffusion values for free AP in buffer and 5 mM ATP (maxima) are 0.72 × 10^−6^ cm^2^/s and 0.97 × 10^−6^ cm^2^/s, respectively which corresponds to an enhancement of 35% (Figure 3B). Whereas the AP tagged vesicle exhibited a maximum diffusion of 0.074 × 10^−6^ cm^2^/s at 5 mM ATP. The increase is ∼23% when compared to the diffusion value of 0.06 × 10^−6^ cm^2^/s for the vesicles in buffer solution (Figure 3C).

**Figure 3:**
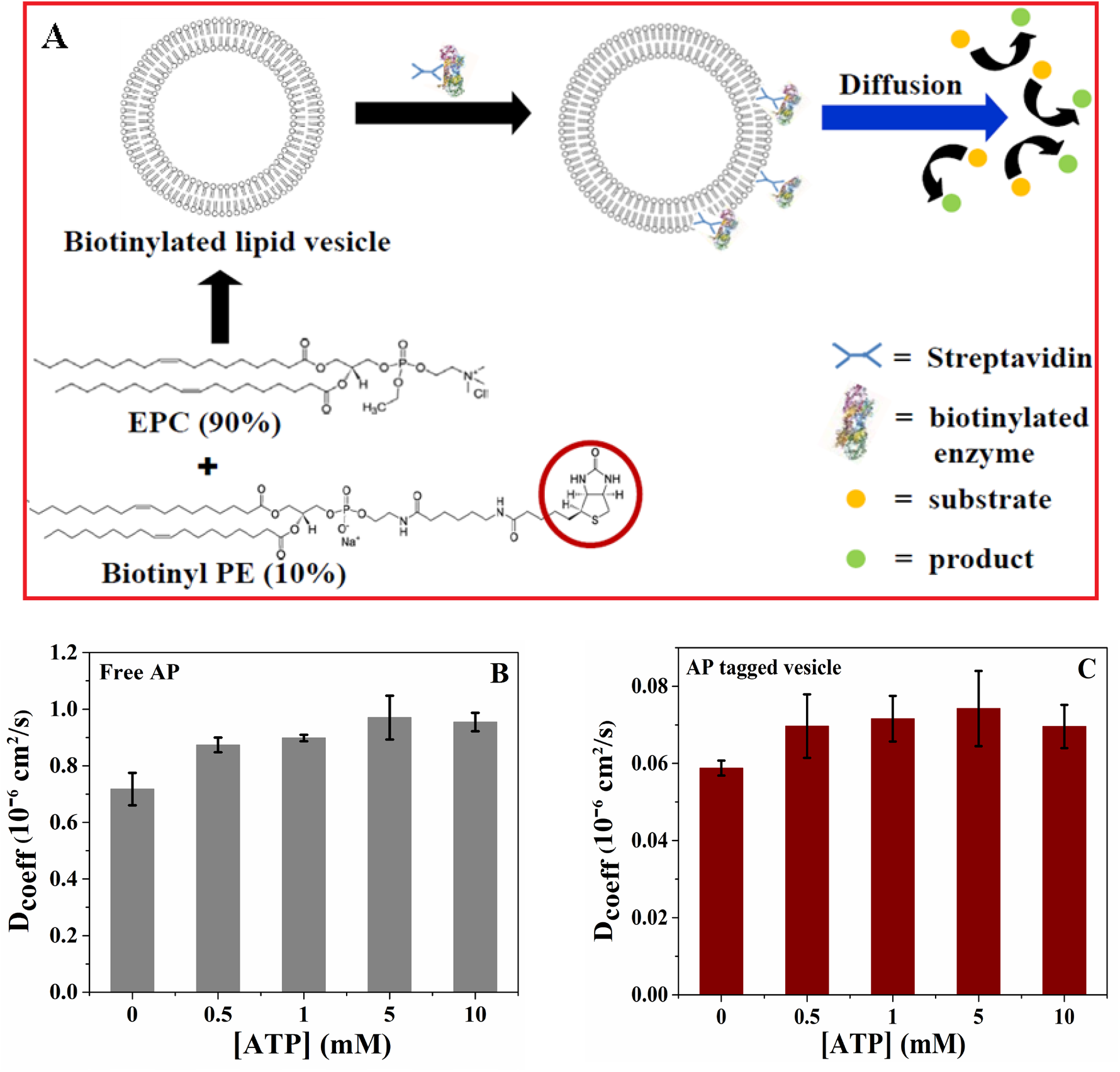
A) Schematic representation showing the lipid composition of the biotinylated vesicles and the subsequent binding of the vesicles to the BS functionalized AP/urease. (B) Diffusion coefficient (D_coeff_) of free AP with increasing ATP concentration. (C) D_coeff_ of AP tagged vesicles with increasing ATP concentration. [ATP] = 0 - 10 mM, [Mg^2+^] = 1 mM. [Citrate buffer] = 10 mM (pH = 5.8) was used to dilute the samples. Diffusion plots were analyzed from the average of five FCS recordings. All the FCS experiments were performed at T=294 K ± 1 K.

FCS analysis of free urease has been well documented to show an enhancement in diffusion in the presence of urea.^36^ In a FCS study of dye labelled free urease, it was observed that diffusion enhancement was 24%, close to the previously reported value of 28%.^36^ The enhancement was estimated from the difference in the diffusion of urease in buffer 0.29 × 10^−6^ cm^2^/s and the diffusion in the presence of 10 mM urea which is 0.36 × 10^−6^ cm^2^/s (Figure S11A). The urease tagged vesicles also exhibited a diffusion enhancement like the other two enzymes ATPase and AP; the diffusion coefficient of urease tagged vesicles was 0.057 × 10^−6^ cm^2^/s in buffer, and 0.070 × 10^−6^ cm^2^/s with 10 mM urea corresponding to an increase of ∼23% (Figure S11B). The diffusion measurements were also repeated for inactive vesicles with varying substrate concentrations and also in the presence of free enzymes in solution. The protocols were similar to the control experiments done for ATPase. In this case, the diffusion coefficient remained unchanged with substrate (Figure S12A,B,C,D).

To further confirm the catalysis-induced increase in diffusion of enzyme attached vesicles, we examined real time motion of catalytically active vesicles in the presence of the substrate using optical microscopy.^37,38^ ATPase bound micron-sized vesicles were mixed thoroughly with different concentrations of ATP and 1 mM Mg^+^ ions and subsequently introduced in a closed cylindrical chamber (details in the Supporting Information). The dimension of the chamber was (D × H) 18 mm × 0.9 mm, and it was carefully fixed to a glass slide before sample addition and subsequently sealed to prevent evaporation and formation of air bubbles, which might generate convective flows in the chamber. The glass slide was placed under an inverted optical microscope and the sample was probed in bright field imaging mode to observe the motion of the vesicles (Figure 4A).^39,40^

**Figure 4:**
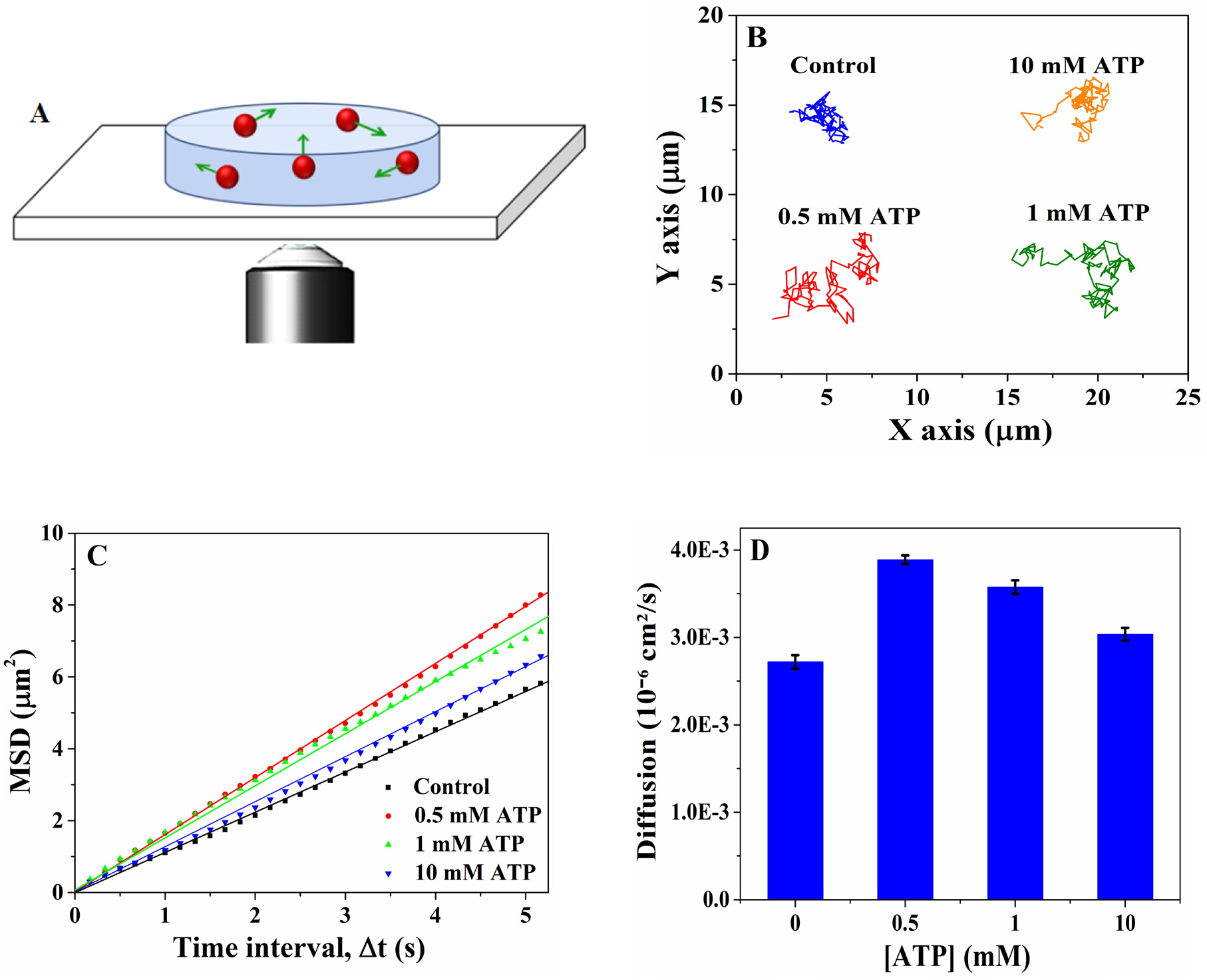
(A) Schematic representation of the hybridization chamber containing ATPase tagged vesicles in a pool of substrate ATP. The movement of the vesicles was observed under the optical microscope at a resolution of 100× and scan rate of 60 frames/seconds. (B) Plot showing a sample trajectory of ATPase tagged vesicles as a function of different ATP concentrations over 20 seconds in the XY plane. Control refers to ATPase tagged vesicles in buffer solution. The trajectories were obtained using the Video Analysis and Modelling Tool. (C) Plots showing the mean square displacement (MSD) of ATPase tagged vesicles for different ATP concentrations as a function of time interval. The motion of at least 10 vesicles was analyzed and averaged for each ATP concentration. (D) Diffusion of ATPase tagged vesicles at different [ATP] as obtained from the slope of the MSDs curves using MSD = 4*D*Δ*t*. The 90% CIs are obtained from the trajectories of at least 10 vesicles followed by curve fitting using Matlab software.

After adjusting the scanning height to 200 μm above the glass surface, video scans were recorded by a high-sensitivity charged-coupled device camera at an optical magnification of 100× and maintaining the scan rate at 60 frames per second for 60 seconds total video time.^40^ After recording the videos, they were analyzed using ‘Tracker’ software to extract the trajectory of the active vesicles and quantify their motion at different ATP concentrations. Figure 4B shows representative trajectories of the ATPase tagged vesicles at different ATP concentrations obtained over 20 seconds. It is clearly evident from the trajectory plots and the videos (Video S1) that the motility of the ATPase tagged vesicles is higher in the presence of ATP, compared to vesicles in only buffer. The motility is highest when the ATP concentration is 0.5 mM, the concentration at which the enzyme has the highest activity as well. On the other hand, the active vesicles remain almost stationary in buffer and moved significantly slower in 10 mM ATP. After extracting the trajectory data from the optical tracking of at least 10 vesicles, mean squared displacements (MSD) of the vesicle ensemble at different time intervals were calculated using MATLAB software and plotted in Figure 4C.^41^ As can be seen, the MSD curves are linear for the tested ATP concentrations. From the slopes of the MSD plot, the effective diffusivity of the vesicles was calculated at each ATP concentration (Figure 4D).

A passive particle immersed in fluid undergoes random diffusive motion as a result of rapidly fluctuating fluid molecules.^42^ An active particle has powered motility in addition to diffusive motion. This powered motility is characterized by steps involving swimming in a specific direction followed by random reorientation.^43^ The characteristic length (*l*) and time scale (*τ*) are, respectively, the step size and the time interval over which there is directional persistence in the particle movement.^44,45^ The active swim velocity of the particle is simply the characteristic length over time scale, *l*/*τ*. For the movement of such active particles, the MSD in 2D as a function of time interval ∆*t* can be written as:

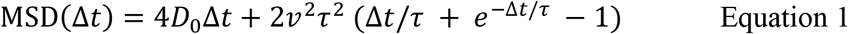

where *D*_0_ is the passive diffusion coefficient of the particle. The first term of the MSD relation is for the diffusive motion and the second part accounts for the powered motility of the particle. The MSD relation can be written at two limiting cases of small (∆*t* ≪ *τ*), and large (∆*t* ≫ *τ*) time intervals as 4*D*_0_∆*t* + 𝑣^2^∆*t*^2^ and 4(*D*_0_ + 𝑣^2^*τ*/2)∆*t*, respectively. At short time intervals, the powered motility of the particle constitutes the ballistic component of its movement (𝑣^2^∆*t*^2^), while at large time intervals, it manifests as an enhancement in the effective diffusion of the particle (𝑣^2^*τ*/2). Therefore, both the ballistic and diffusive regimes are the result of the powered motility at different observation time scales, ∆*t*. The observed enhancement in the diffusion (Figure 4D) is an indication of powered motility of the vesicles in the presence of ATP.

The trend of diffusion enhancement of active vesicles obtained from optical microscopy is similar to that observed by FCS and both correlate well with the enzyme activity at different ATP concentration (Figure 2C). Thus, the optical tracking results along with the FCS measurements further establish active motion of the enzyme-bound vesicles upon catalytic turnover of their respective substrate.

## Conclusion

The diffusive motility of ATPase tagged vesicles in the absence and presence of substrate ATP was studied by FCS. The diffusion coefficient of the active vesicles was found to increase with increasing enzymatic turnover rate. Using different triphosphates, UTP, CTP and GTP, we showed that the vesicle motility is closely correlated with the ATPase activity for each of these substrates as well. Similar behavior was also observed for vesicles with either AP or urease attached to the vesicle wall via biotin-streptavidin linkage. The powered motility of the ATPase tagged vesicles was also substantiated in real time by optical microscopy tracking experiments and MSD analyses. The diffusion data from optical microscopy experiments closely resemble the diffusion data from FCS. Thus, the observed enhanced diffusion is unlikely due to enzyme dissociation into subunits or due to FCS related artefacts.^20^ Our results constitute the first steps in the fabrication of biocompatible, multifunctional hybrid motors for carrying out specific functions under physiological conditions. The membrane-bound protocells that move by transducing chemical energy into mechanical motion also serve as models for motile living cells and are key to the elucidation of the fundamental mechanisms governing active membrane dynamics and cellular movement. Future studies to quantify the enzymatic forces that impart self-generated motion and parallel investigations on the mechanical fluctuations in vesicle membranes are in progress.

## Acknowledgement

The work was supported by the Center for Chemical Innovation funded by the National Science Foundation (CHE-1740630). FM acknowledges support from Penn State MRSEC (DMR-1420620)

